# A synoptic revision of *Pohliella* (Podostemaceae) with notes on *Aulea, Cipoia* and *Saxicolella*

**DOI:** 10.1101/2020.05.23.111922

**Authors:** Martin Cheek

**Affiliations:** Herbarium, Royal Botanic Gardens, Kew, Richmond, Surrey, U.K

**Keywords:** Amphi-atlantic, Conservation, Extinct, Phylogeny, Rheophyte

## Abstract

*Saxicolella* Engl., an African genus of the waterfall specialist plant family Podostemaceae, was shown to be polyphyletic as currently delimited. One clade, sampled from species in Ghana, is sister to American *Ceratolacis* (Tul.)Wedd., *Podostemum* Michx. and all Old World Podostemoideae (podostemoids). The second clade, sampled from Cameroonian material, was embedded within the major clade of African podostemoids. In this paper the generic nomenclature applied to *Saxicolella* sensu lato (*Saxicolella, Pohliella* Engl.*, Aulea* Lebrun & Stork nom. inval.), is reviewed and the morphological support for the two clades and their correct generic names is determined. *Pohliella* is shown to be the correct name for the first clade (based on *Pohliella laciniata* Engl., Cameroon) and a synoptic treatment of its three published species is given, one of which is extinct, and two are threatened. However, a fourth, unpublished species exists. The new combinations *Pohliella submersa* (J.B.Hall) Cheek and *Pohliella amicorum* (J.B. Hall) Cheek are made for the two published Ghanaian species. The recently described New World genus *Cipoia* C.T. Philbrick, Novelo & Irgang is revealed as being morphologically identical to *Pohliella*, but in view of the geographical disjunction, confirmation from molecular evidence is awaited before its two species are also transferred to *Pohliella*. The correct name for the second clade, embedded in African podostemoids, is *Saxicolella* (*sensu stricto*), now with two known species, *Saxicolella nana* Engl. (type of *Saxicolella,* Cameroon) and *Saxicolella flabellata* (G.Taylor) C. Cusset (Nigeria).

## INTRODUCTION

Podostemaceae are a pantropical family of annual or perennial herbs placed in Malpighiales in a sister relationship with Hypericaceae (*Ruhfel et al. 2011*). There are about 300 species globally, in c. 54 genera (*Koi et al. 2012*). Species numbers are highest in tropical America, followed by Asia, with Africa having c. 106 species (*Cheek & Lebbie 2018*). All species of the family are restricted to rocks in rapids and waterfalls of clear-water rivers (rheophytes) or occur in the spray zones of waterfalls. However, waterfalls are being increasingly exploited for hydropower at some risk to the survival of the Podostemaceae they contain (*Schenk et al. 2015, Cheek et al. 2015, Cheek & Ameka 2016, Cheek et al. 2017b*). Most of the African species of Podostemaceae are narrow endemics, many being known from only a single waterfall. New discoveries of species are still made frequently (*Rial 2002, Cheek 2003, Schenk & Thomas 2004, Beentje, 2005, Cheek & Ameka 2008, Kita et al. 2008, Cheek et al. 2015, Schenk et al. 2015, Cheek & Ameka 2016, Cheek & Haba 2016, Cheek et al. 2017a, Cheek et al. 2017b, Cheek & Lebbie 2018, Cheek et al. 2019, Cheek et al. 2020*).

Three subfamilies are recognised. Tristichoideae, sister to the rest of the family, have three foliose tepals that protect the developing flower, and are tricarpellate. Weddellinioideae, with a single genus are Neotropical and have two foliose tepals and bilocular ovaries. Podostemoideae, pantropical, is the most genera and species rich subfamily. It has flowers protected in a spathellum, a balloon-like sac in which the flower develops while the plant is underwater, and tepals reduced to vestigial, filiform structures. African Podostemoideae, or podostemoids, are the main focus of this paper.

Important characters in defining genera in African podostemoids are the position of the flower in the unruptured spathellum, and the number of locules, shape, and sculpture of the ovary. At species level, important characters are the shape and relative proportions of spathellae, stigmas, anthers, filaments, gynophores, pedicels, and leaves.

The current taxonomic framework for African Podostemaceae was set in place by the revisions and Flora accounts of Cusset (*Cusset 1973, 1974, 1978, 1983, 1984, 1987 and 1997*). Only recently has accumulating molecular phylogenetic data begun to influence the classification (*Moline et al. 2007, Thiv et al. 2009, Schenk et al. 2015)* Cusset’s work has been compiled and updated by *Rutishauser et al.* (*2004*) who recognise c. 85 species in 16 genera.

However, *Saxicolella* Engl. was one of the few genera that Cusset did not revise. Yet, in her Flore du Cameroun account (*Cusset, 1987*), in addition to the type species *Saxicolella nana* Engl., she included in *Saxicolella* the genus *Pohliella* Engl. with two species *P. laciniata* Engl. and *P. flabellata* G. Tayl. *Taylor (1953)* had already expressed his doubts about *Pohliella* “Apart from differences in habit and shape of the stigmas, I am not satisfied that the key characters used by Engler to distinguish *Pohliella* from *Saxicolella* are sufficiently diagnostic.” Despite this doubt, after equivocating, Taylor described *Pohliella flabellata* from Nigeria. In the next paragraph he described a new genus, also from Nigeria, *Butumia* G. Tayl. “most closely related to *Saxicolella and Pohliella* “. *Butumia* was transferred to the first genus as *Saxicolella marginalis* (G. Tayl.) Cheek having been discovered in Cameroon (*Cheek et al. 2000:153*). *J.B. Hall (1971),* published three new Podostemaceae, all endemic to Ghana, including *Saxicolella amicorum* J. B. Hall. A second of his species, described as *Polypleurum submersum* J.B. Hall was later also transferred to *Saxicolella* by *Cook & Rutishauser (2001).* All these species share a flower which is erect in the spathellum, remaining included within it at anthesis, with a single stamen and dyad pollen. Ameka *et al.* (2002) presenting the developmental morphology of *Saxicolella amicorum* and *S. submersa* (J.B.Hall) C.D.Cook & Rutish. from Ghana, recognised six species of *Saxicolella* and stated it to be the third largest genus among the African Podostemaceae. In confirming the bilocular ovaries of *S. amicorum* and *S. submersum* that had first been reported by *J.B Hall (1971), Ameka et al. (2002)* speculated that “this may have been the reason why *Cusset (1987)* may have segregated the obviously bilocular species *S. amicorum* and *S. submersum* in the provisional genus *Aulea* which was never officially published (see Lebrun & Stork, 1991)”. They further stated that “In our view locule number alone should not be a reason for creating a new genus for *S. amicorum* and *S. submersum*.” *Ameka et al. (2002)* then justified at length their caution regarding locule number as a generic character.

The name *Aulea* first appears in print in *Lebrun & Stork (1991)* including two species: “*A. amicorum* (J.B. Hall) C. Cusset, inédit and *A. submersa* (J.B. Hall) C. Cusset, inédit”. However *Aulea* is not validly published according to the Code (*Turland et al. 2018*), so it and the binomials which include it are *nomina nuda*. Cusset herself is not known to have published on the subject of *Aulea*.

The African Plant Database (continuously updated), which is based on the work of Lebrun & Stork (e.g. *Lebrun & Stork, 1991*), gives *Aulea amicorum* as deriving from an unpublished manuscript. Cusset was based at National Museum of Natural History, Paris (P). Two specimens retrieved as *Aulea* at P are both identified in the hand of C. Cusset as *Aulea amicorum* (MNHN collections website (continuously updated)). These are 1) *J.B. Hall* in GC 37132 from the Ankasa River in Ghana, a paratype of *S. amicorum* J.B. Hall, and 2) *Aké Assi* 11649 ‘Dans la lit de la Rivière Tabou à Yalea, Cote D’Ivoire. The last appears to be an *Inversodicraea* on the basis of its scale-leaves, a synapomorphy of that genus in Podostemoideae. These determinations suggest that either Cusset’s concept of *Aulea amicorum* is based on heterogenous elements, or that one of her determinavit slips was glued to the wrong sheet.

The generic name *Aulea,* attributed to C. Cusset by *Lebrun & Stork (1991)*, but never validly published, took on a life of its own.

*Cook & Rutishauser (2001)* in discussing *Saxicolella* and in formally transferring to this genus *Polypleurum submersum* J.B. Hall, state that “Cusset in *Lebrun & Stork (1991)* assigned *S. amicorum* and *S. submersa* to a supposed new genus “*Aulea*”, which has never validly been published. We assume she considered the presumptive unilocular ovary to be sufficient to distinguish “*Aulea*” from *Saxicolella*, which has a bilocular ovary. Ameka and Rutishauser (unpubl.) have found that both *S. amicorum* and *S. submersa* have bilocular ovaries, which negates the basis of “*Aulea*”.” *Cook & Rutishauser (2007)* and *Rutishauser et al (2004)* then treat *Aulea* as a synonym of *Saxicolella.* Yet the statement above is incorrect in that Cusset herself never published this assignment, nor is there a basis for presuming that she considered *Saxicolella* to have bilocular ovaries. In fact, in her only published work that included *Saxicolella* she stated that it has unilocular ovaries (*Cusset 1987:92*). She would have been aware of *Hall’s (1971)* depiction of the ovary of the Ghanaian taxa as bilocular. Notably, *Cusset (1987)* did not include Ghana in the range of *Saxicolella,* consistent with the idea that she saw the Ghanaian material as a separate genus that she provisionally named *Aulea* (writing determination slips with this name), but which for unknown reason she did not see fit to publish.

The most complete molecular phylogenetic overview of the Podostemaceae to date (*Koi et al. 2012*) includes three species of *Saxicolella*, the type species *S. nana* Engl., and two Ghanaian species, *S. amicorum,* and an undescribed species. The resulting tree (*Koi et al. 2012*: Fig. 1) shows the type species *S. nana* to be embedded in a clade comprising almost all the African Podostemoideae, with the exception of the two Ghanaian species of *Saxicolella.* These two species form a strongly supported second African clade which is shown as sister (weakly supported) to a more inclusive clade comprising the Podostemoideae of all other African, Asian and Australian Podostemoids, plus two American genera, *Podostemum* A. Michx. and *Ceratolacis* (Tul.) Wedd.

**Fig. 1.**
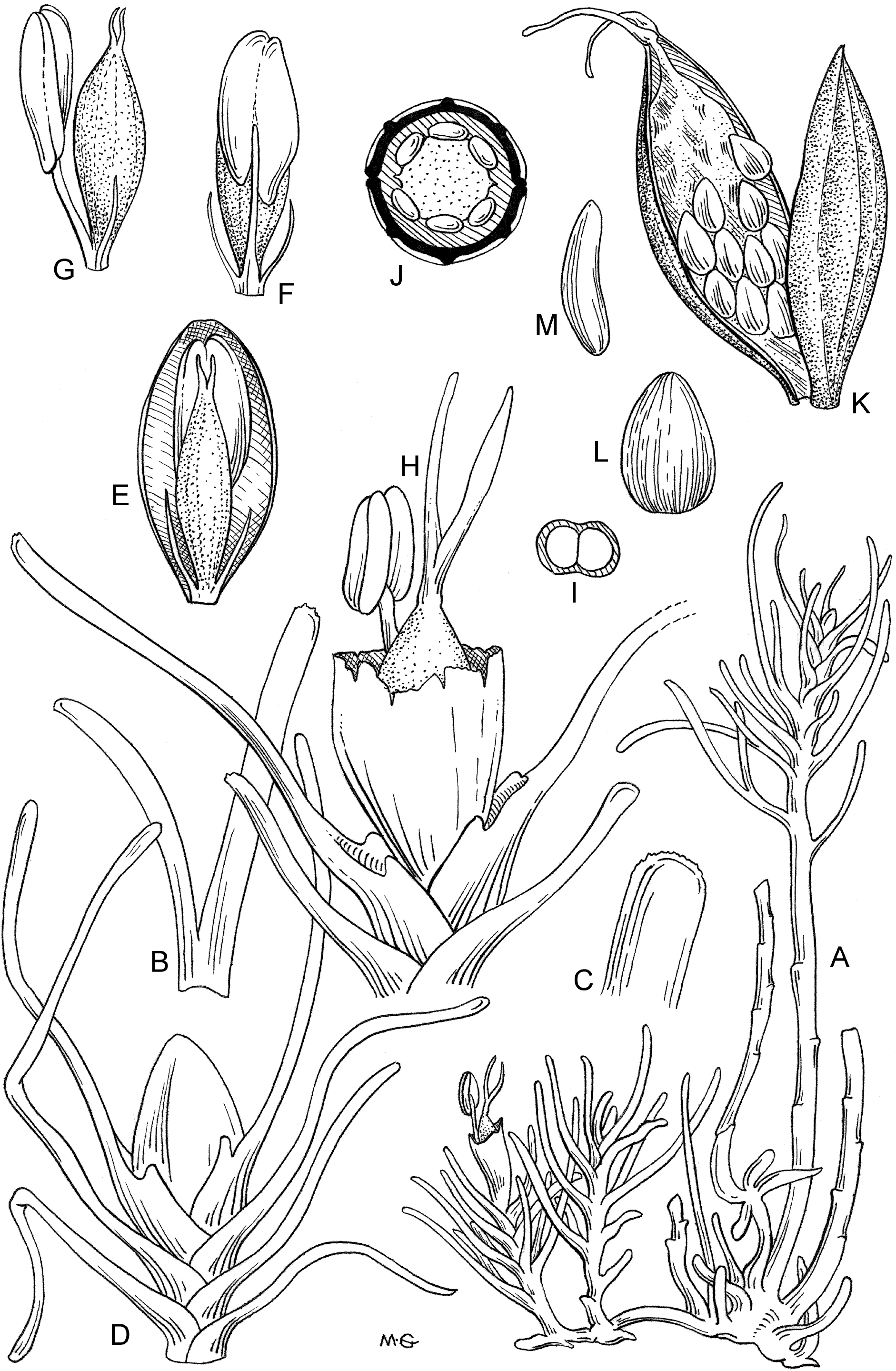
Pohliella amicorum. (J.B. Hall) Cheek reproduced from Hall (1971) with permission.

Sister to that large clade plus Ghanaian *Saxicolella,* are the rest of American Podostemoid genera apart from *Diamantina* Novelo, C.T. Philbrick & Irgang, suggesting an American origin for Podostemoideae. Therefore, it appears from reading *Koi et al. (2012),* that, from an origin in S. America, the Podostemoideae crossed the Atlantic to West Africa before crossing back to the Americas (*Podostemum* and *Ceratolacis*) and then, again recrossing the Atlantic back to Africa and the rest of the Palaeotropics.

Thus *Saxicolella*, following the concept of *Cook & Rutishauser (2001), (2007)* and *Rutishauser et al. (2004)* is polyphyletic, with two distinct clades *(Koi et al. 2012).*

The objective of this paper is to determine whether there is morphological support for the two clades of *Saxicolella*, and if so, what are their correct generic names and included species, focussing on the clade that is sister to the rest of the Old World podostemoids.

## MATERIAL AND METHODS

Nomenclatural changes were made according to the Code *(Turland et al., 2018).* Names of species and authors follow IPNI (continuously updated). Observations in Ghana were made as part of the Rheophytes of Ghana project (*Ameka et al. 1996*). Regarding the Cameroonian species, this study was part of a series of botanical surveys, mainly led by the author, which began in 1991. The methodology used is as reported in *Cheek & Cable (1997)*, and specimen data storage by Gosline, in *Cheek et al.* (*2004*). Fieldwork approvals, permits and agreements are as reported in *Cheek et al.* (2018).

The morphological species concept was followed in defining species (each species being separated from its congeners by several, usually qualitative, morphological disjunctions), and the overall morphology of species was described and illustrated following standard botanical procedures as documented in *Davis & Heywood (1963).* All specimens cited have been seen by the author unless indicated ‘n.v.’ Herbarium citations follow Index Herbariorum *(Thiers et al., continuously updated*) and authors of plant names *IPNI* (*continuously updated*). Material was studied from B, BM, EA, GC, HNG, K, L, P, SCA and YA.

Conservation assessments were either taken from the recent literature (see citations) or made using the categories and criteria of *IUCN (2012).* The cell-size used for calculating area of occupancy was 4 km^2^, as advocated by IUCN. Herbarium material was examined with a Leica Wild M8 dissecting binocular microscope. This was fitted with an eyepiece graticule measuring in units of 0.025 mm at maximum magnification.

## TAXONOMIC TREATMENT

There is considerable morphological support for recognition of two clades within what was hitherto known as *Saxicolella.* Two character-states that separate the clades are given by *Koi et al. (2012),* citing *Ameka et al.* (*2002*). A further three character-states are given in this paper. The five character states that separate the two clades (see table 1 below) are more than sufficient to merit generic separation.

**Table 1.** Characters separating the two clades of *Saxicolella*

|  | *Saxicolella* clade sister to Old World Podostemoideae ('Ghanaian *Saxicolella*', or *Pohliella*) | *Saxicolella* clade embedded in African Podostemoideae (*Saxicolella* s.s.) | Data source |
| --- | --- | --- | --- |
| Root caps | Present* | Absent | Ameka *et al.* 2002 |
| Phyllotaxy of main stems | Distichous** | Spiral | This paper |
| Root morphology | Subfiliform (slightly dorsiventrally flattened), <1 mm width | Broadly ribbon-like (width >10 mm) or crustose | This paper |
| Paired root holdfasts | Present | Absent | This paper |
| Ovary | Bilocular | Unilocular | Ameka *et al.* 2002 |

It remains only to determine the correct names for the two genera. *Saxicolella* based on *S. nana* is the earliest name for the genus embedded in the main, large African clade of Podostemoideae. The earliest generic name for the sister to all the Old World Podostemoideae is *Pohliella*, based on *Pohliella laciniata* Engl. of Cameroon, which is congeneric in terms of the characters in table 1, with the Ghanaian species formerly attributed to *Saxicolella* by *Cook & Rutishauser (2001, 2007)* and *Rutishauser et al.* (*2004*) and to *Aulea* by *Lebrun & Stork (1991)* and by *Koi et al. (2012).*

Thus *Engler (1926)* who first published these two genera was, after all, correct in his original delimitation, although not supported by any subsequent author, until the present. However, as pointed out by *Taylor (1953),* the key characters used by Engler to distinguish *Pohliella* from *Saxicolella* (e.g. rib number and stigma shape) are insufficiently diagnostic. *Saxicolella* sensu stricto will be the subject of a future taxonomic synoptic revision. Here is presented a synoptic revision of the genus *Pohliella*, sister to all Old World podostemoids on the evidence of *Koi et al. (2012)*.

### Synopsis of *Pohliella* Engl

***Pohliella*** Engl. (*Engler 1926*:457) non *Taylor* (*1953:53a; 1954*)

Type species: *Pohliella laciniata* Engl.

*Saxicolella* pro parte sensu auctt. e.g. *Cusset (1987)* quoad *S. laciniata* (Engl.) Cusset, sensu *Hall (1971)* quoad *S. amicorum* J.B. Hall, non Engl. (*Engler 1926*).

*Polypleurum* sensu *J.B. Hall (1971)* quoad *P. submersum* J.B. Hall, non (Tul.) Warm. (*Warming 1901*).

*Aulea* Lebrun & Stork nom. inval. (*Lebrun & Stork 1991:79*).

*Cipoia* C.T. Philbrick, Novelo & Irgang (*Philbrick et al. 2004*)

<u>Rheophytic herbs</u>. Roots terete, thread-like, or slightly dorsiventrally flattened (to about twice as wide as thick), 0.2–0.9 mm wide, adhering to the substrate by root hairs on the ventral surface and by haptera. Haptera exogenous, lateral to the roots, in pairs, disc-like, or comprised of one or more thread-like strands. <u>Stems</u> endogenous, arising above the haptera, erect, terete, 5–50 cm long, 0.4–0.8 cm diam., unbranched, or rarely branched. <u>Leaves</u> alternate, distichous, bases sheathing, with a pair of acute basal stipules in leaves subtending spathellae, blades canaliculate at base, flat, ribbonlike, or filiform, entire or with 1 or more bifurcations, rarely trifurcate, apices acute. <u>Flowers</u> single, terminal on short axillary shoots, or on main stems, or in clusters terminal on main stems. <u>Spathellae</u> ellipsoid, sessile, apices rounded. <u>Flowers</u> erect in bud, not inverted, at anthesis mostly or partly concealed inside the spathellae, pedicels accrescent in fruit; tepals 2, filiform, erect. Stamen1, inserted just above tepals. Pollen in dyads. Gynophore short. Gynoecium narrowly ellipsoid, 2-locular. Stigmas 2, filiform, erect. Capsule with 8 longitudinal ribs. Seeds numerous, ellipsoid, mucilaginous when wetted.

**Phenology:** flowering in early dry season as water-levels drop

**Distribution & habitat:** four species in Ghana and Cameroon, of which three are formally named (the fourth is the undescribed species from eastern Ghana of which I have seen no material: ‘*S. agumatsa* Ameka & Cheek’ fide *Koi et al. (2012)* based on *Ameka* 478 (G.C. n.v.), 479 (GC n.v.)); rapids in evergreen forest or in woodland. There appear to be two additional species in Minas Gerais, Brazil which are currently placed in *Cipoia* (see discussion below). The Cameroonian species occurs in the most species-diverse area of Cameroon (*Cheek et al. 2001*), while the Ghanaian species are scattered from west to east, each in a separate vegetation type.

The ecology of *Pohliella* as observed for *P. amicorum* in Ghana and *P. laciniata* in Cameroon by the author, is different from that of most other African podostemoids (Cheek pers. obs. 1995–2018). Both these *Pohliella* species occur in rivers partly shaded by lowland evergreen rainforest and where, even in in the dry season, the water-levels remain high, as already observed by *Hall (1971)* for *P. amicorum*.

Unlike the majority of African Podostemaceae, with notable exceptions such as *Ledermanniella letouzeyi* (*Cheek et al. 2004*), those species of *Pohliella* observed (see above) seem to persist, and to spread underwater vegetatively by their thread-like roots, forming colonies that can be tens of metres long, rather than, as in most other African podostemoids, patches confined to waterfalls or rapids in full sun which dry out and die as annuals in the dry season (Cheek pers. obs. 1992–2018).

**Etymology:** named for Joseph Pohl (1864–1939) of Wroclaw, foremost botanical artist over 40 years for the prolific German botanist Adolf Engler *(Engler 1926).* Pohl produced many tens of thousands of botanical drawings, including more than 33 000 drawings in 6 000 plates for just one of Engler’s larger works *(Anon 2018).*

**Local names:** none are known.

**Conservation:** all three described species are threatened (see species accounts below).

*Notes* – Detailed micromorphological, developmental, palynological and anatomical studies of *Pohliella* (as *Saxicolella amicorum and S. submersum*) are depicted in Ameka et al. (2002) which is a key reference to *Pohliella,* since it treats in great detail two of the four species of the genus. Nothing is known of the chromosomes, chemistry, germination, or of the pollination or reproductive biology of *Pohliella.* In comparison with most African podostemoids, such as *Ledermanniella* and *Inversodicraea,* the flowers of *Pohliella* are extremely inconspicuous. In the case of *P. laciniata,* only the two dull, red filiform styles and the white anther emerge from the green spathellum at anthesis, and often only a single flower per inflorescence is open at a time. In fact, it was not obvious that the plants were in flower (Cheek pers. obs. 2008). Nevertheless, insect pollination is most likely as has been observed for other African podostemoids (Cheek et al. 2017). However, *Ameka et al. (2002)* speculate that wind pollination occurs in *P. amicorum,* and Hall (1971) that water pollination might be possible in *P. submersa.*

*Koi et al. (2012)* indicated that the systematic affinities of *Pohliella* (referred to as *Saxicolella* or *Aulea* of Ghana) are with Podostemoideae in America. In that molecular phylogenetic analysis, *Pohliella* is sister to *Podostemum* and those two genera together are sister to *Ceratolacis*. The two *Pohliella* taxa sampled in that analysis were *Pohliella amicorum* based on *Ameka & DeGraft-Johnson 112, 113* (GC n.v.) and the undescribed *Pohliella* referred to as ‘*Saxicolella agumatsa* Ameka & Cheek’ (see above and *Koi et al. 2012*)

*Philbrick et al. (2004)* describe two new American genera. *Diamantina* is sister to all Podostemoideae and does not concern us here (*Koi et al. 2012*). *Cipoia* C.T. Philbrick, Novelo & Irgang however, is placed as close to *Podostemum* and *Ceratolacis*, the three being characterised as the dyad group since their pollen grains are dispersed in pairs. All other American podostemoids have monads (single pollen grains), except for *Diamantina* which has tetrads (*Philbrick et al. 2004*). *Cipoia,* now with two species (*Bove et al.2006*), is noteworthy because there are no points of separation morphologically between it and *Pohliella* (Cheek pers. obs.). All of the features that characterise *Cipoia* (*Philbrick et al. 2004*) are found in *Pohliella* as described here, e.g. the single stamen per flower, the ovary and mature capsule remaining enclosed within the ruptured spathellum during and after anthesis, with only the stigmas and stamens projecting from the spathellum. Equally *Cipoia*, like *Pohliella*, has thread-like roots, a gynophore and isolobous, blunt-apexed, bilocular ovary-fruit.

Were it not for the fact that the two groups of species are separated by the Atlantic, and that no other case is known of amphi-atlantic genera in the podostemoids (although *Tristicha* in Tristichoideae is amphi-atlantic), I would have no hesitation in immediately uniting and transferring the two species of *Cipoia* to *Pohliella.* I propose that material of *Cipoia* be sequenced and, if as expected, a sister relationship with *Pohliella* is evidenced, to then unite them. *Soltis et al. (2007)* give another example of a long overlooked amphi-atlantic sister relationship.

### Identification key to the published species of *Pohliella*

1. Flowers single or in clusters; leaves of non-flowering stems flattened, blade-like, laciniate. Cameroon **1. *Pohliella laciniata*** Flowers single; leaves of non-flowering stems more or less cylindrical, entire or forked. Ghana 2
2. Stems 1.5-12 cm tall, self-supporting, ± unbranched, flowers terminal; in shade of evergreen forest. Extant. **2. *Pohliella amicorum*** Stems 25 cm long, flowing in water, flowers terminal on numerous short side-shoots; open woodland in full sunlight. Extinct. **3. *Pohliella submersa***

#### 1. Pohliella laciniata

Engl. (*Engler 1926*:457; *1930*:49 fig. 39).

Holotype: Cameroon, Nkongsamba, Bakaka Forest, Near Babong, Dinger River “steine und felsen im reissenden” (stones and rocks in the torrent), 400 m a.s.l., fl.fr. 21 Nov. 1908, *Ledermann* 1185 (holotype B, B100160608!)

> *Saxicolella laciniata* (Engl.) C. Cusset (*Cusset 1987*:94; Cheek in *Onana & Cheek 2011*: 253, 504c) Homotypic synonym.
>
> *Inversodicraea laciniata* Engl. (*Engler 1915*:274, fig. 177) Homotypic synonym.

**Phenology:** flowering and fruiting in November and December.

**Distribution & habitat:** endemic to SW Region Cameroon; torrents in lowland evergreen forest areas; in the Mone River this species was locally the most abundant of five species of Podostemaceae present together, the others being *Tristicha trifaria* (Willd.) Spreng., *Inversodicraea ledermannii* Engl. (*Cheek* 14352), *Ledermanniella* cf. *variabilis* (*Cheek* 14354A) and *Ledermanniella* cf. *linearifolia Cheek* 14351). When observed at Mone on 8 Dec. 2008, the first two species had already long been exposed by the falling river level, dried and fruited, while the last two species, occurring deeper in the water and more recently exposed, were actively flowering and fruiting, as was *Pohliella laciniata.* At Korup on the Mana River, the specimen was collected along with other rheophytes: *Macropodiella pellucida* (Engl.)C.Cusset (2652), *Inversodicraea* sp. (2653), Podostemaceae indet. (2655), *Kanahia laniflora* (Forssk.)R.Br. (2656), *Habenaria weileriana* Schltr. (2657), *Phyllanthus dusenii* Hutch. (2658) and *Virectaria angustifolia* (Hiern)Bremek. (2659); 50–500 m alt.

**Etymology:** referring to the laciniate leaves.

**Additional specimens:** Cameroon, SW Region, 20 km W Mamfe, path from Agborkem to Tabo, River Bawan, st. June, *Letouzey* 13731 (P n.v.); Korup National Park, 4°58”N, 8° 51”W, fl.fr. 9 Dec. 1983, *D. Thomas* 2654 (K, MO n.v., P n.v.); Mone Forest Reserve, Mone River downstream from Amebishu, fl.fr. 8 Dec. 2008, *Cheek* 14350 (K!, SCA, YA).

**Conservation:** *Pohliella laciniata* (as *Saxicolella laciniata*) was assessed as Endangered, EN B1+B2ab(iii) since three locations were known (Cheek in *Onana & Cheek, 2011*). It was considered possibly extinct at the type location and to face threats due to siltation from run-off resulting from palm oil plantations and illegal logging documented at the other two. Extent of occurrence was calculated as 3006 km^2^ and area of occupation 3 km^2^ using 1 km^2^ cells. This assessment is maintained here since no new data are available in the last ten years, excepting that a fourth location is now known. The additional location results from the redetermination of *Thomas* 2654 (K) which had previously been identified as *Saxicolella flabellata* (G.Taylor) C.Cusset (*Cusset 1987*). This species appears genuinely rare and range-restricted since while Cameroon is not exhaustively studied for Podostemaceae, it is well studied in comparison with neighbouring countries in Africa. Most of the African samples included in previous global molecular phylogenetic studies of Podostemaceae have derived from Cameroon (*Ruhfel et al. 2011, Koi et al. 2012*).

### Notes

The slender, far-running, black, thread-like roots are anchored at regularly spaced nodes by opposite groups of three radiating thread-like holdfasts (Cheek field obs. 2008). Clusters of subsessile spathellae develop above each set of holdfasts. From these clusters arise erect, unbranched (rarely once-branched from base) slender stems 1–2 cm long with alternate, distichous, subentire broad leaves. The stems terminate in a leafy, cabbage-like rosette (*Cheek* 14350, K).

*Pohliella laciniata* has the largest range of any species of the genus. It occurs, or occurred, in three different catchment areas: the rivers Dinger (type location), tributary of the Wouri; the Bawan and the Mone, tributaries of the Cross; and the Mana which flows directly into the sea. This area corresponds with the Cross-Sanaga interval, which has the highest species diversity per degree square in tropical Africa (*Cheek et al. 2001*) including numerous other threatened, endemic species, such as *Kupea martinetugei* Cheek (*Cheek et al. 2003*), *Ancistrocladus grandiflorus* Cheek (*Cheek 2000*) and *Cola metallica* Cheek (*Cheek 2002*).

#### 2. Pohliella amicorum

(J.B. Hall) Cheek comb. nov.

Holotype: Ghana, Ankasa F.R., forming dense cover on rocks just above and below low watermark. Sterile below surface; fertile above. in the river bed, fl. fr. 31 Dec. 1966. *Hall, Enti & Jenik* in GC 36241 (K00419997 holotype!, K000042000 isotype!, GC isotype!) Fig. 1

> Basionym: *Saxicolella amicorum* J.B. Hall (*Hall 1971:133*).

**Phenology:** flowering and fruiting in November and December.

**Distribution & habitat:** Ghana, Ankasa River, river torrents in lowland evergreen forest, “forming dense cover on rocks just above and below low watermark. Sterile below surface; fertile above” (*Hall et al*. 36241, K).

**Etymology:** commemorating the friendship which the co-collectors (Enti, Jenik and John Hall) of the type of this plant enjoyed during their excursions through the forests of SW Ghana (*Hall 1971*).

**Additional specimens:** Ghana, Ankasa R., on rocks in the river bed, fl. fr. 11 Nov. 1967, *Hall* in GC 37132 (GC!, K000041998!, K000041999!, P00179135!); Western Region, Ankasa Resource Reserve, Ankasa river, 21 Nov. 1998, *Ameka 113* (GC n.v.).

**Conservation:** *Pohliella amicorum* is known only from a short section of the Ankasa River so that one event upstream could threaten the entire global population at the several sites downstream. Therefore it is here treated as having a single location in the sense of *IUCN* (*2012*). The area of occupancy is estimated as <20 km^2^ With projected threats from surface run-off upstream at risk of contaminating the river with silt and nutrients (which are death to Podostemaceae), the species is assessed here as Critically Endangered, CR B1+B2ab(iii).

*Notes - Pohliella amicorum* is unusual among African Podostemaceae in growing in shaded locations (pers. obs. Cheek 1995).

#### 3. Pohliella submersa

(J.B. Hall) Cheek comb. nov.

Holotype: Ghana, Kwahu-Nteso, rapids of river Asuboni, fl. fr. 17 Jan 1968, *Hall & Bowling* in GC 38532 (K holotype; GC isotype), Fig. 2

**Fig. 2.**
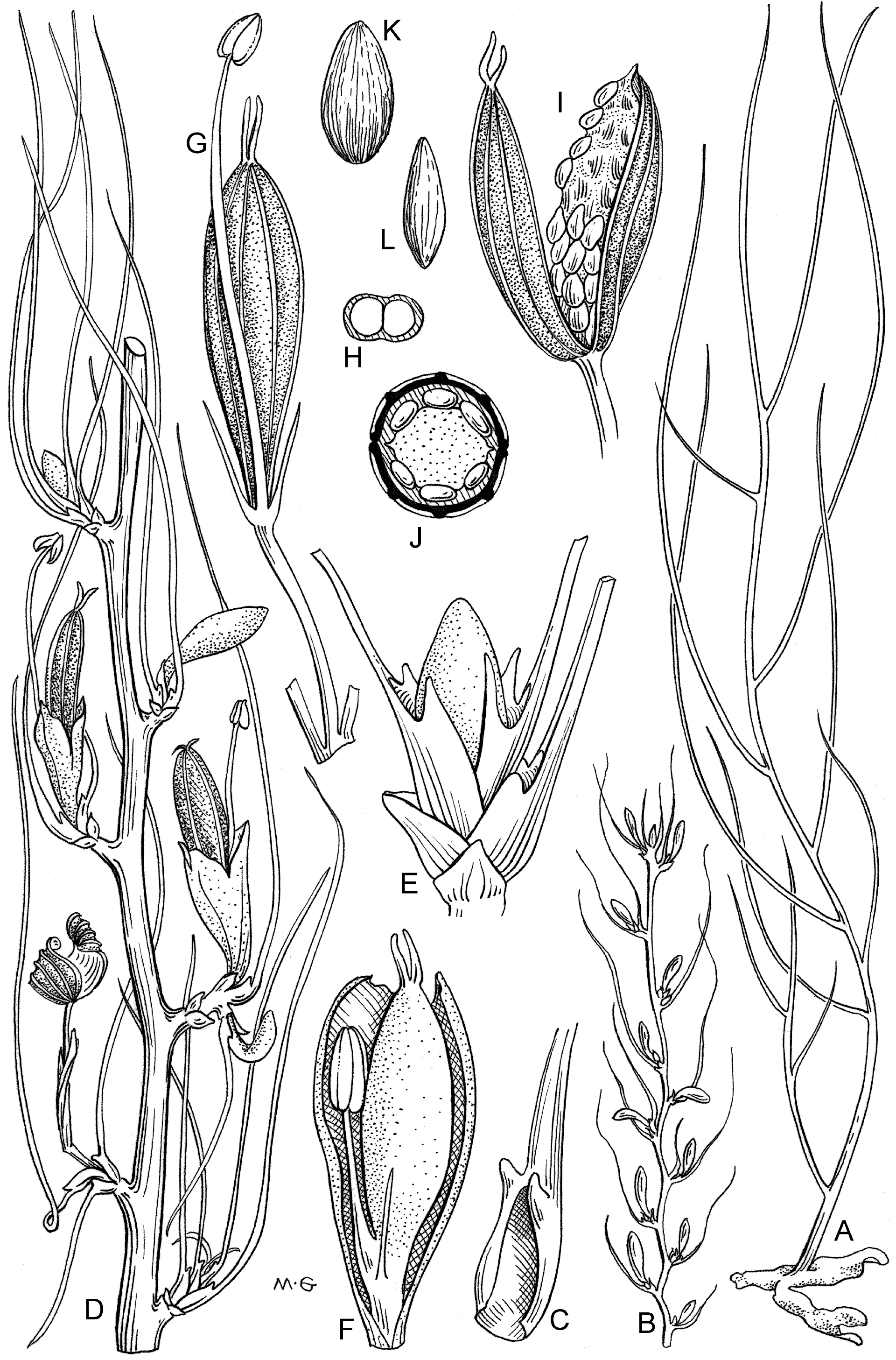
Pohliella submersum. (J.B. Hall) Cheek reproduced from Hall (1971) with permission.

Basionym: *Polypleurum submersum* J.B. Hall (*Hall 1971*:130)

*Saxicolella submersa* (J.B. Hall) C.D.K.Cook & R. Rutish. (*Cook & Rutishauser 2001*:1165). Homotypic synonym.

**Phenology:** flowering and fruiting in January.

**Distribution & habitat:** torrents in lowland dry forest areas

**Etymology:** referring to the flowing, submersed stems.

**Additional specimens:** R.Akrum nr Begoro, “reddish straggling plant forming submerged bushes in slow moving pool 18” deep”, st. 12 June 1970, *JB Hall* in GC 39649 (K!); ibid fl. 5 Feb. 1971, *JB Hall* in GC 42400 (K!);

**Conservation:** known only from the two collection sites cited above. Targetted efforts to refind this species beginning 30 years after its discovery have failed (Ameka and Cheek pers. obs). This species is now thought to be extinct (*Ameka et al. 2002*). However, there is a remote possibility that it may yet be found at some as yet unsearched rapids. Accordingly it is here assessed as CR B1+B2ab(iii), i.e. Critically Endangered (Possibly Extinct) since it was known from a two locations with threats due to nitrification of the water - algal growth on the plants as noted in the protologue.

Notes: this species was intensely investigated by Ameka, together with *P. amicorum*, for his PhD thesis (*Ameka 2000*), later published as *Ameka et al.* (*2002*). Since no live plants could be found, he used spirit preserved material at K for his micro-morphological studies. *Hall* (*1971*) mistakenly gave the root width as 8 mm, but *Ameka et al.* (*2002*) indicated that the correct measurement is 0.2–0.5 mm.

## EXCLUDED SPECIES

*Pohliella flabellata* G. Taylor (*Taylor 1953*:53) is referable to *Saxicolella flabellata* (G. Taylor) C. Cusset.since it has a foliose root and a unilocular ovary (*Taylor 1953*:53).

## DISCUSSION

The dyad group of podostemoid genera arose in the Americas in a sister relationship with the monad group (*Koi et al.2012*). It is hypothesised here that *Pohliella* is basal to the dyad group and migrated from America (where currently named as *Cipoia*) to Africa.

The *Pohliella* clade is not widespread nor species-rich in Africa. One can speculate that this may be due to morphological and physiological restrictions that limit its colonisation or reduce its competitiveness in the waterfalls now dominated by the second migration of podostemoids from America, of which the most basal branch in Africa is *Inversodicraea* (*Koi et al.2012*). However, many African genera, for want of material, were not included in the molecular phylogenetic study of *Koi et al.* (*2012*). When these gaps are filled other genera are likely to contend with *Inversodicraea* for the basal branch position, such as *Stonesia* G. Taylor (excluding the morphologically and geographically discordant *Stonesia ghoguei* E.Pfeifer & Rutish. which was the only *Stonesia* included in *Koi et al.2012*) and especially *Lebbiea* Cheek (*Cheek & Lebbie, 2018*). *Ceratolacis* in America, sister in the latest study (*Koi et al. 2012*) to the major African Podostemoid clade resulting from the second migration already shows what may be a key morphological innovation that may have contributed to the success of the second podostemoid migration to Africa which unlike the first resulted in a major radiation of genera and species. This innovation is a widening of the root which can reach 5 mm wide in *Ceratolacis* (*Philbrick et al. 2004*, *Cook & Rutishauser, 2007*). A broader root offers the possibility of a stronger purchase on the substrate, allowing plants with this feature to better colonise white-water conditions which may disadvantage more basal podostemoids which have thread-like roots <1 mm wide. Additional innovations arising in Africa that may have contributed to the major radiation are reproductive:

1. Loss of the ovary septum allowing more complete opening of the capsule and dispersal of seed.
2. Inversion of the flowers in the spathellum, allowing development in bud of a long pedicel permitting the flowers to extend both further and more rapidly above the stems and the water surface. This innovation, seen in the most successful (the most geographically widespread and species-diverse) genera *Inversodicraea* and *Ledermanniella*, can be speculated to improve pollination success, by removing the flowers quickly from the pollen-killing effects of water and by making them more accessible to winged insect pollinators such as honey bees. In American dyad species the pedicel is only 2–3 mm long (measurements made from illustrations in *Philbrick & Novelo 2004*) while in the African radiation pedicel length is longer, 7–8(–15) mm long in *Inversodicraea* (*Cheek et al. 2017b*).

## CONCLUSIONS

If the hypothesis presented in this paper is to be tested, that *Pohliella* as here defined, including *Cipoia,* is sister to the rest of Old World podostemoids together with *Podostemum* and *Ceratolacis*, then improved generic-level sampling, ideally with good species-level representation, are needed for incorporation in molecular phylogenetic studies.

## ACKNOWLEDGEMENTS

Janis Shillito typed the manuscript. Eimear Nic Lughadha and two anonymous reviewers gave advice on earlier versions of the manuscript. Mike Swaine (University of Aberdeen) is thanked for developing the Rheophytes of Ghana project through the Darwin Initiative and, with Gabriel Ameka, James Adomako, Diggy de Graft-Johnson, thanked for companionship on visits to Ghana. Aaron Nicholas and Ymke Warren (Wildlife Conservation Society, Cameroon) supported fieldwork to the Mone Forest Reserve in Cameroon, and Messrs E. Ndive (Limbe Botanic Garden), Sama (WCS), J Tampie, D. Tampie, T. Tako and M. Obi (of Mukojong and Mbu villages) provided assistance as field botanists, assistants and guides. Julia Buckley of the Department of Library, Art and Archives, Royal Botanic Gardens, Kew is thanked for arranging with the estate of Mary Grierson for permission to re-use Figures 1 and 2, which were originally labelled as *Saxicolella amicorum* and *Polypleurum submersum* and appeared in *Hall(1971).*

